## Supplemental Tables for "CD4 T cell Responses in the CNS during SIV infection"

**S1 Table. Nonhuman primate cohort for current study.**

| **Group** | **Animal ID** | **Sex** | **Age (years. months) at Nx.** | **Weight (kg) at Nx.** | **Site** | **Virus**  **(1x10^4^ TCID_50_)** | **IgG antibody* control (25mg/kg)** | **ART**  **Regimen** |
| --- | --- | --- | --- | --- | --- | --- | --- | --- |
| Acute 251 Cohort (n=4) | 41812 | F | 11.02 | 7.30 | CNPRC | SIVmac251 | Anti-desipramine | - |
|  | 40742 | M | 12.04 | 12.00 | CNPRC | SIVmac251 | Anti-desipramine | - |
|  | 37164 | F | 16.01 | 10.83 | CNPRC | SIVmac251 | Anti-desipramine | - |
|  | 35488 | F | 17.04 | 10.48 | CNPRC | SIVmac251 | Anti-desipramine | - |
| Chronic 251 Cohort (n=6) | 38889 | F | 14.06 | 11.31 | CNPRC | SIVmac251 | - | FTC/TDF/DTG |
|  | 38919 | F | 14.06 | 8.63 | CNPRC | SIVmac251 | - | FTC/TDF/DTG |
|  | 36056 | F | 18.06 | 9.96 | CNPRC | SIVmac251 | - | FTC/TDF/DTG |
|  | 37274 | F | 16.06 | 8.67 | CNPRC | SIVmac251 | - | FTC/TDF/DTG |
|  | 39359 | M | 14.04 | 13.36 | CNPRC | SIVmac251 | - | FTC/TDF/DTG |
|  | 36511 | F | 17.06 | 10.34 | CNPRC | SIVmac251 | - | FTC/TDF/DTG |
| Controls (n=4) | 38163 | F | 15.07 | 7.48 | CNPRC | - | - | - |
|  | 40691 | M | 12.07 | 12.18 | CNPRC | - | - | - |
|  | 38691 | F | 14.10 | 13.56 | CNPRC | - | - | - |
|  | 40499 | F | 12.11 | 9.79 | CNPRC | - | - | - |

SIV: Simian Immunodeficiency Virus; ART: Anti-Retroviral Therapy; FTC: Emtricitabine; TDF: Tenofovir disoproxil fumarate; DTG: Dolutegravir; CNPRC: California National Primate Research Center

*****Antibody treatments were administered at 1 week prior to SIV infection and every subsequent 10-day interval at doses of 25mg/kg intravenously (neat) over the course of the study.

**S2 Table. Antibody reagents for Flow Cytometry Analysis.**

| **S. No** | **Reagents** | **Source** | **Catalog number** |
| --- | --- | --- | --- |
| 1. | AF700 anti-human CD3 (Clone SP34-2) | BD Biosciences | Cat# 557917 |
| 2. | APC-Cy7 anti-human CD3 (Clone SP34-2) | BD Biosciences | Cat#557757 |
| 3. | BV650 anti-human CD4 (Clone L200) | BD Biosciences | Cat# 563737 |
| 4. | BUV805 anti-human CD8 (Clone SK1) | BD Biosciences | Cat#612889 |
| 5. | BUV737 anti-human CD95 (Clone DX2) | BD Biosciences | Cat# 564710 |
| 6. | PE/Dazzle™ 594 anti-human CD28 (Clone CD28.2) | BioLegend | Cat# 302942 |
| 7. | APC-Cy7 anti-human CD20 (Clone 2H7) | BioLegend | Cat#302314 |
| 8. | APC-Cy7 anti-human live/dead | invitrogen | Ref#L34976A |
| 9. | PE anti-human CD197 (CCR7) (Clone 3D12) | BD Biosciences | Cat# 561008 |
| 10. | BV785 anti-human CD195 (CCR5) (Clone 3A9) | BD Biosciences | Cat#565001 |
| 11. | PE-CF594 Mouse Anti-Human CD196 (CCR6) (Clone 11A9) | BD Biosciences | Cat# 564816 |
| 12. | BV711 anti-human CD69 (Clone FN50) | BioLegend | Cat#310944 |
| 13. | FITC anti-human CD49d | Beckman Coulter | Part no# IM1404U |
| 14. | PE/Dazzle™ 594 anti-human/mouse Integrin β7 (Clone FIB504) | BioLegend | Cat#321226 |
| 15. | PE anti-human Integrin β1 (Clone TS2/16) | BioLegend | Cat# 303003 |
| 16. | APC anti-human CD183 (CXCR3) (1C6/CXCR3) | BD Biosciences | Cat#550967 |
| 17. | PECy7 anti-human PD1 (Clone EH12.2H8) | BioLegend | Cat# 329918 |
| 18. | FITC anti-human TNF-α (Clone Mab11) | BioLegend | Cat# 502906 |
| 19. | PECy7 anti-human IFNγ (Clone B27) | BioLegend | Cat# 506518 |
| 20. | PE/Dazzle™ 594 anti-human IL-2 (Clone MO1-17H12) | BioLegend | Cat# 500344 |
| 21. | FACS lyse | BD Biosciences | Cat#349202 |
| 22. | FoxP3/ Transcription Factor Staining Buffer set | invitrogen | Cat#00-5523 |
| 23. | Brilliant stain buffer | BD Biosciences | Cat#563794 |

CD: cluster of differentiation; PE: Phycoerythrin; AF: Alexa Fluor; BV: Brilliant violet; BUV: Brilliant ultraviolet; Cy: Cyanine; APC: Allophycocyanin; FITC: Fluorescein Isothiocyanate Conjugate.
