## Supplemental figures for "CD4 T cell Responses in the CNS during SIV infection"

**S1 Figure. CCR5/CCR7 dichotomy during homeostasis/Blood counts during acute SIVmac251 infection**

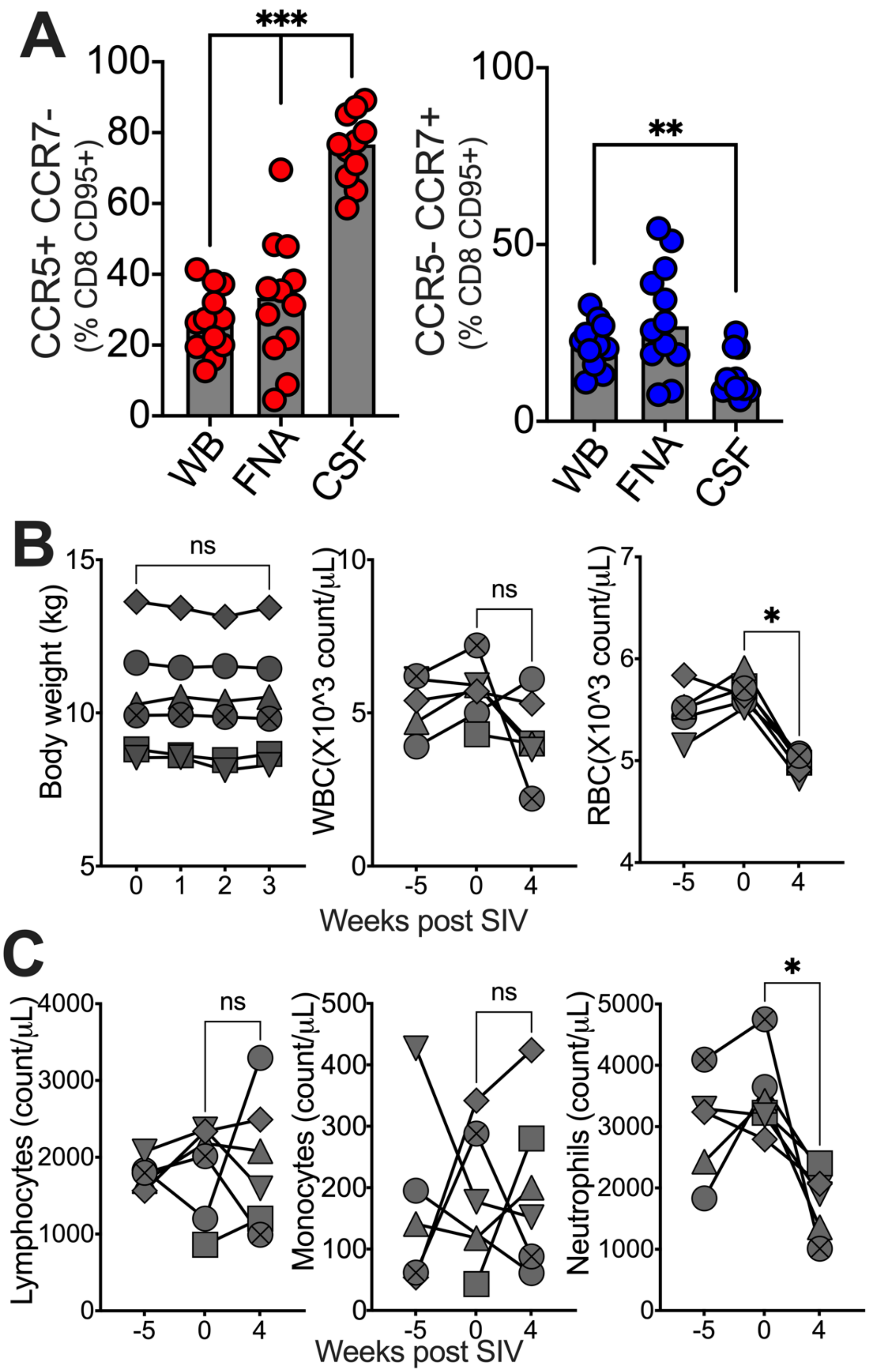

**S2 Figure. Plasma and CSF cytokines during acute SIVmac251 infection**

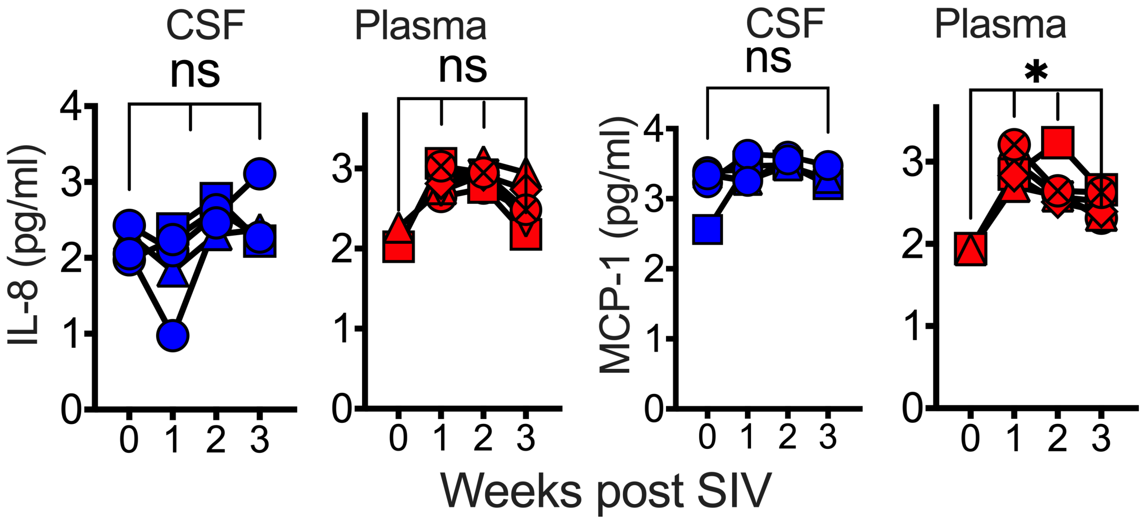


**S3 Figure. CSF CCR5+ CD8 T cell frequencies do not decrease during acute SIV infection**

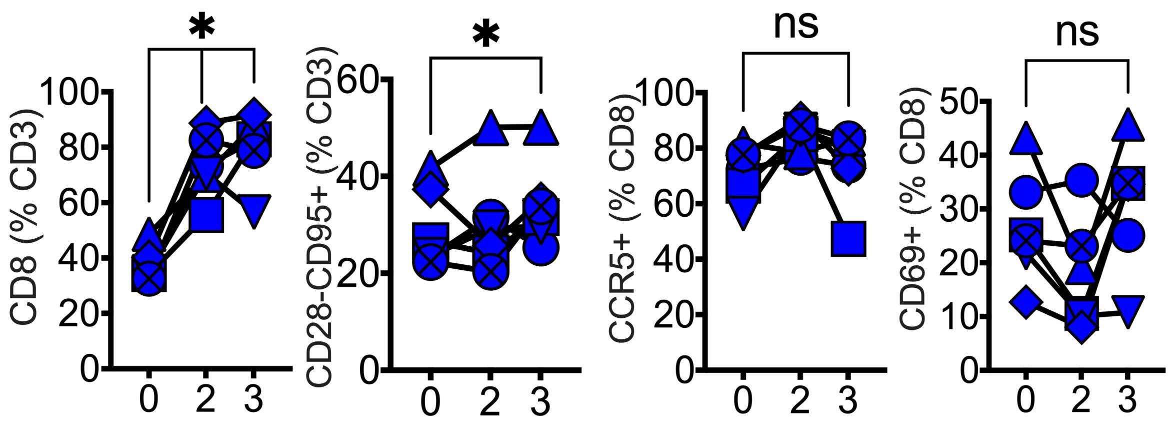

**S4 Figure. CSF parameters during acute SIVmac251 infection**

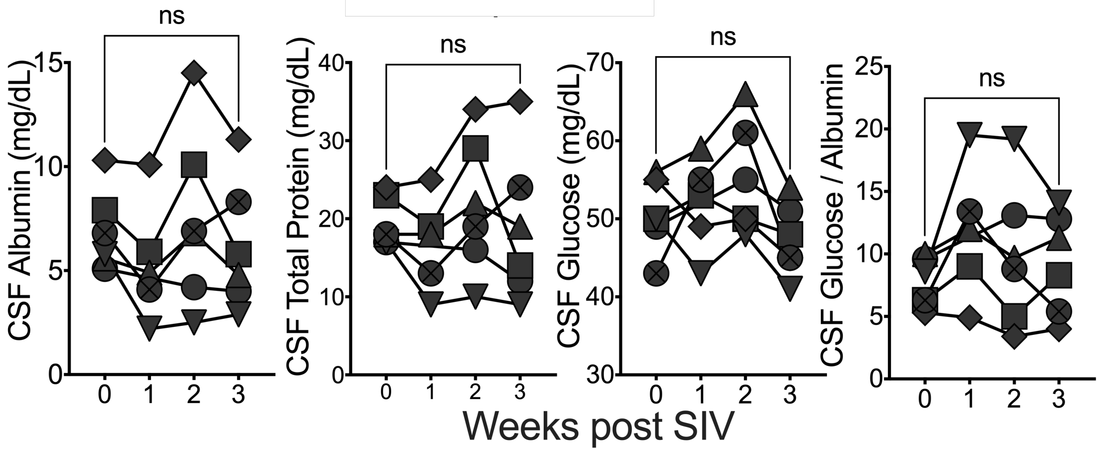


**S5 Figure. CCR5+ CD4 T cells populate parenchymal and border CNS tissues**

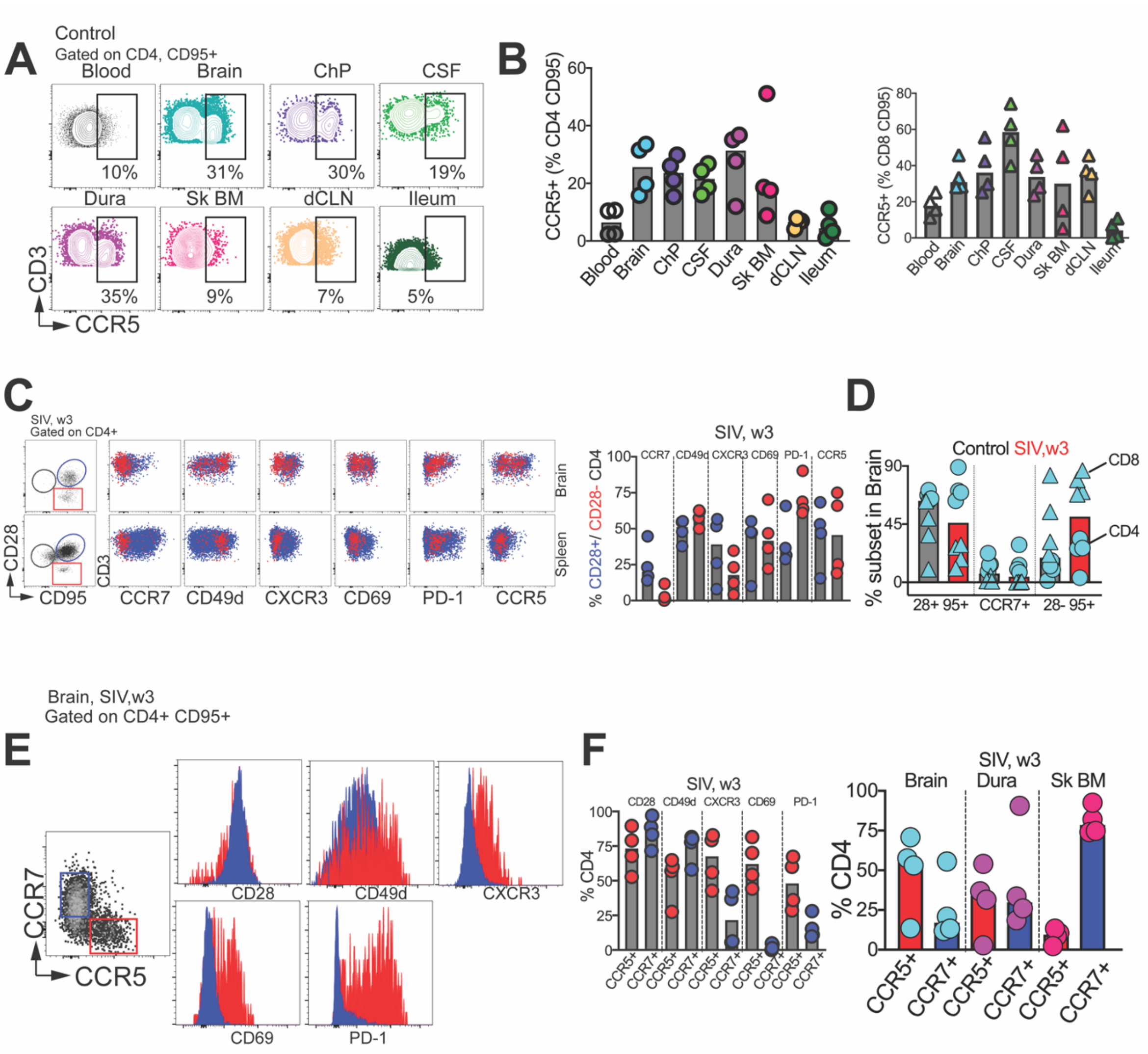

### S6 Figure. Volcano plots of DEG in SIV brain

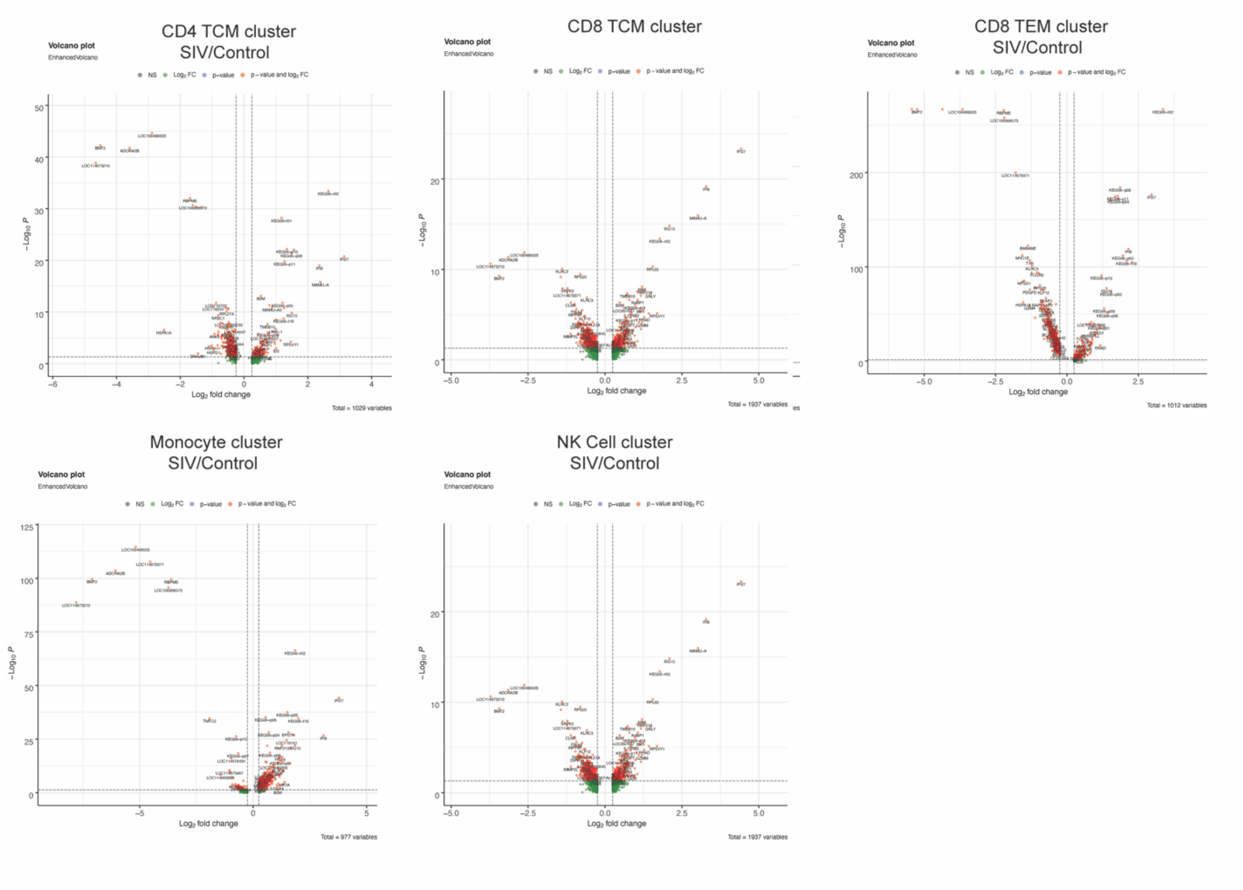

**S7 Figure. Pseudo time plots of Control and SIV brain**

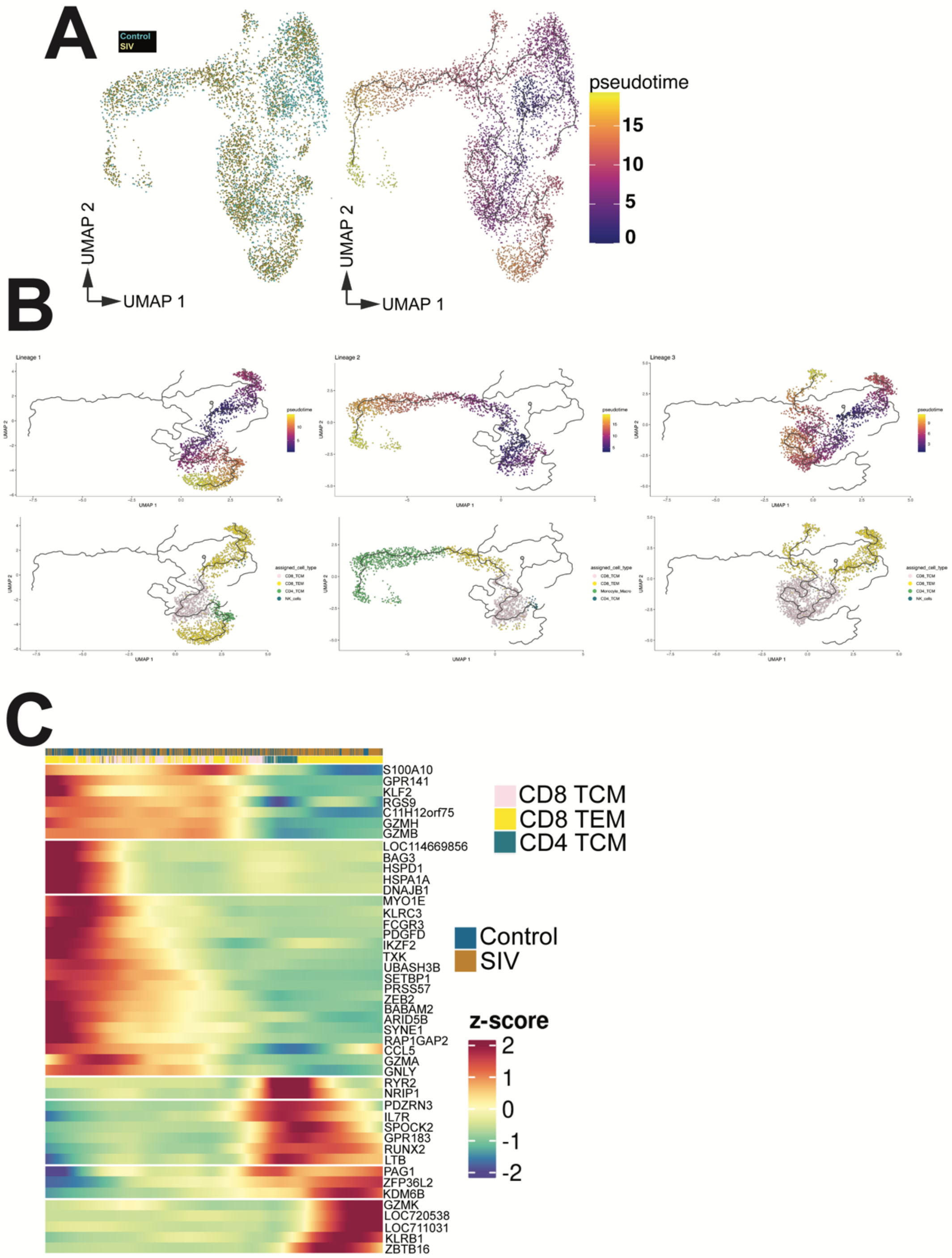

#### S8 Figure. Kinetics of plasma and CSF vRNA, plasma IP-10 and CSF IP-10

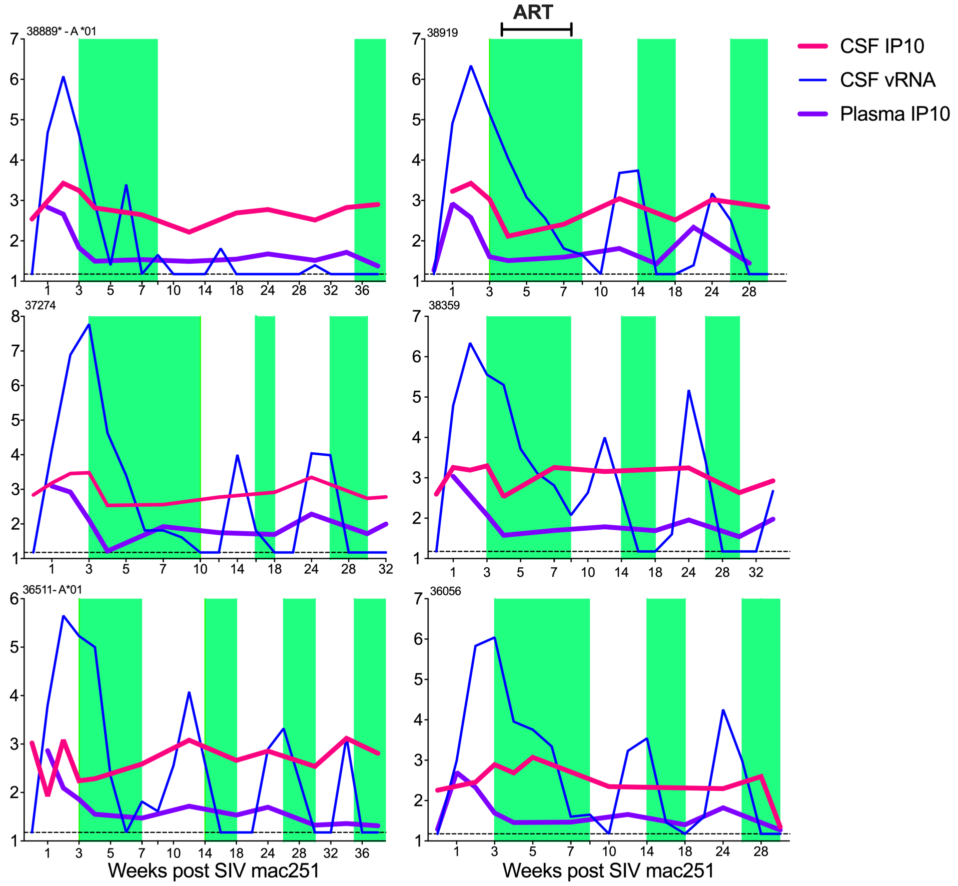
